# Geodesic theory of long association fibers guidance on the human fetal cortex

**DOI:** 10.1101/2023.01.29.526048

**Authors:** Kazuya Horibe, Gentaro Taga, Koichi Fujimoto

## Abstract

Association fibers connect different areas of the cerebral cortex over long distances and integrate information to achieve higher brain functions, particularly in humans. Prototyped association fibers are developed to the respective tangential direction throughout the cerebral hemispheres in the subplate layer during the fetal period. However, the directional guidance principle for forming association fibers is unknown. Because the subplate is located below the cortical surface, the tangential direction of the fibers may be biased by the curved surface geometry due to Sylvian fissure and cortical poles. The fiber length can be minimized if the tracts follow the shortest paths (geodesics) of the curved cortical surface. Here, we propose and examine a theory that geodesics guide the tangential direction of long association fibers by analyzing how geodesics are spatially distributed on the fetal human brains. Unlike the homogeneous distribution on spherical surfaces, we found that the geodesics were dense on the saddle-shaped surface of the perisylvian region and sparse on the dome-shaped cortical poles. The geodesics corresponded with the positions of five typical association fibers, supporting the geodesic theory. Thus, the geodesic theory provides directional guidance and suggests that long association fibers emerge from minimizing their tangential length on fetal brains.

## Introduction

Association fibers achieve higher brain functions in humans and connect long-distance special anatomical areas of the cerebral cortex to integrate their functions (Philippi et al., 2009; Egger et al., 2015; Mori et al., 2005; Catani et al., 2012). There are five major association fibers, which connect Broca’s area to Wernicke’s area (the arcuate fasciculus), the cingulate gyrus to the entorhinal cortex (the cingulum (cg)), the frontal pole to the temporal pole (the uncinate fasciculus (unc)), the frontal pole to the occipital pole (the inferior fronto-occipital fasciculus (ifo)), and the occipital pole to the temporal pole (the inferior longitudinal fasciculus (ilf)), based on Mori’s atlas (Mori et al., 2005). Association fibers begin to form early in the fetal period at around 13 gestational weeks (GW), and all major fibers emerge by 20 GW (Horgos et al., 2020; Takahashi et al., 2012; Huang et al., 2009). Although the wiring of interhemispheric commissural fiber on the coronal section is well understood at the molecular level, including the axon guidance (Comer et al., 2019; Stoeckli, 2018), a method to wire the long association fibers in a tangential direction within the hemisphere has not yet been discovered, likely due to experimental difficulty in cutting out the respective brain sections.

Association fibers are initially formed at the subplate layer, which appears only during the fetal period directly below the cortical layer (Kostovic and Rakic, 1990; Kanold and Luhmann, 2010; Duque et al., 2016; Hadders-Algra, 2018; Molnár et al., 2019; Vasung et al., 2019). The subplate layer is a tangentially continuous tissue throughout the cerebral hemispheres (Kostovic and Rakic, 1990) and is well-developed in primates, especially in humans (Vasung et al., 2016). Subplate neurons are sparsely distributed in this layer and are surrounded by glial cells and extracellular matrix, making the subplate layer uniform and elastic (Kostović et al., 1989). Subplate neurons are locally connected to each other through gap junctions and chemical synapses (Moore et al., 2014; Kristt and Molliver, 1976; Luhmann and Khazipov, 2018; Luhmann et al., 2022). The locally connected subplate neurons show the membrane potential activities spontaneously or stimulus-responsively (Molnár et al., 2020; Moore et al., 2011; Ohtaka-Maruyama et al., 2018; Ohtaka-Maruyama, 2020) which can be propagated tangentially throughout the layer (Kilb et al., 2011; Garaschuk et al., 2000; Sun and Luhmann, 2007). Moreover, despite the immaturity of the cortical plate at approximately 24 GW, electroencephalogram (EEG) and functional magnetic resonance imaging (fMRI) findings have indicated the presence of global neuronal activities throughout the cerebral hemispheres, likely derived from the subplate (Thomason et al., 2017; Keunen et al., 2017). Therefore, the subplate nexus hypothesis has been proposed to explain the developmental mechanism of association fibers. According to this hypothesis, the subplate layer, as a uniform continuum tangentially stretched across the cerebral hemispheres, creates the long-distance, “weak” neuronal fibers (precursor of association fibers) by locally connecting neuronal fibers during the fetal period (Kostović, 2020).

Because the subplate in the fetal period is located below the cortical surface (Kostovic and Rakic, 1990), fiber formation may be biased by the smoothly curved landscape in the form of domes at the brain poles with positive Gaussian curvature and saddles on the perisylvian region (appearing around 11–29 GW) with negative Gaussian curvature (Fig. 1, left panel) (Kostović et al., 2019; Van Essen, 2020). Fiber formation between a pair of distant positions on the cortical surface has a higher cost, as the structural materials of the neuronal fibers (neurofilaments, microtubules, etc.) must cover a longer tract (Kanold and Luhmann, 2010; Kostović et al., 2019). Therefore, wiring likely follows the shortest tract to minimize the costs (Fig. 1, middle panel). These assumptions allow us to derive a hypothetical directional guidance principle for the long-distance fibers, hereafter referred to as the geodesic theory, in which the shortest path (i.e., geodesic (Kobayashi, Shoshichi and Nomizu, 1963) of a cortical surface provides the tangential direction of the prototyped association fibers.

**Fig. 1:**
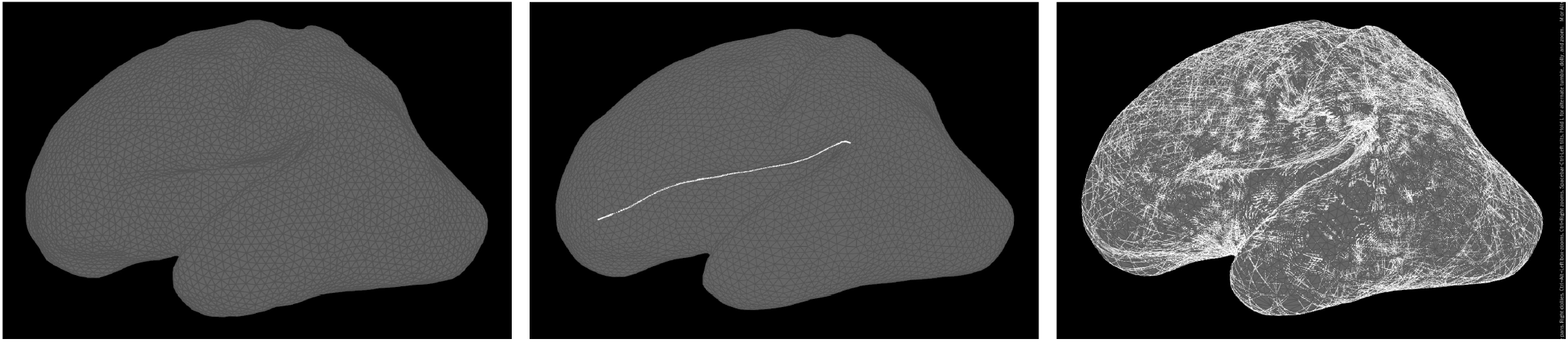
Representative methods used in this study. At 27 GW the cortical surface geometry was represented by triangular meshes (left), and a geodesic between a pair of randomly selected positions was calculated (white line: a single geodesic, center). Overwriting 1, 500 geodesics covered the entire brain surface (right)

The direction of the geodesics generally depends on the surface curvature (Kobayashi, Shoshichi and Nomizu, 1963). Geodesics, each connecting two arbitrary positions on the surface, are uniformly distributed on a sphere with spatially uniform curvature and can be non-uniformly distributed on surfaces with non-uniform curvature (e.g., sparse at domes with positive curvature and dense at saddles with negative curvature). Similarly, the brain poles and fissures/sulci are dome- and saddle-shaped, respectively. However, the spatial distribution of geodesics on cortical surfaces lacks evidence (Fig. 1, right panel). Assuming that a single geodesic corresponds to a single neuronal fiber, a bundle of multiple geodesics may account for the association fibers. Comparing the spatial distribution of geodesics with those of long-association fibers could validate the geodesic theory.

An appropriate stage for this comparison is around 27 GW, because at this time, the subplate layer comprises a large volume and all major association fiber prototypes are accumulated within the subplate layer (Kostović, 2020). At this age, the Sylvian fissure impact the overall landscape of the smooth surface of the subplate layer and the cortical layer, prior to the gyrification of the primary and secondary sulci (Van Essen, 2020). Hence, in this study, we analyzed the cortical surface geometry at 27 GW to validate the geodesic theory.

## Materials and Methods

### Brain samples

We used the left brain surface data of a preterm infant with 27 GW published by Garcia et al. (Garcia et al., 2018b). The corpus callosum was cut out, and the cross-section was masked. The brain surface data in *gii* format downloaded from BALSA (https://balsa.wustl.edu/) were converted to *obj* format files using the GIfTI library of Matlab (MathWorks Inc., USA) to create triangular mesh data. To modify the edge length of the triangular mesh to a mean of 1.25 mm, we sequentially applied a re-meshing function (uniform method, iterations = 3, smoothing = 0.1) and a smoothing function (curvature-dominant method) in Houdini (version 18.5; SideFX Inc., Canada) to the mesh. Then, the mesh of the corpus callosum was removed using Houdini’s Blast geometry node, resulting in 9,998 triangular mesh faces. The codes are available on our GitHub page (https://github.com/KazuyaHoribe/GeodesicTheory).

### Geometrical analysis

We calculated the shortest paths (geodesics) on the cortical surface using the eikonal equation:

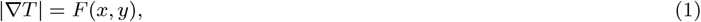

where *T* is the distance function and *F* (*x, y*) is the velocity of movement of the boundary. In this study, we set *F* (*x, y*) = 1. To obtain a finite element approximation of the eikonal equation, we used the fast marching method (Kimmel and Sethian, 1998; Sethian, 1996), which provides a distance function on the cortical surface between the start and end positions, which were sampled randomly among the nodes of the triangular meshes. Using *T*, the geodesic to the initial position was obtained by solving the following differential equation (Eq. 2, Fig. 1, central panel).

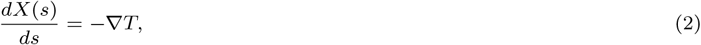

where *X*(*s*) represents the geodesic and *ds* = 0.625 mm (distance of half of the edge length). From the end position, we sequentially computed the curve on the surface following the gradient of the distance function *T* while completing the termination condition as follows.

The distance from the starting position was less than *ds* = 0.625 mm or the inner product of the movement vector was negative, i.e., the direction of movement was reversed. Thus, the geodesic was given by a sequence of points and a complementary curve between them.

We calculated 50,000 geodesics on the cerebral cortex and confirmed that the ensemble of the geodesics covered almost the entire surface area. As an indicator of the local density of the geodesics, we computed the passage frequency of the geodesics at each node of the mesh. If the position of points representing the geodesic was contained within a radius of 0.5 mm (nearly half of the edge length) of each node, the geodesic was judged to pass through that node. The Gaussian curvature on the mesh was calculated using the Gaussian function in Houdini.

We visualized the correspondence of the geodesics to the association fibers by applying “OR” and “AND” operations. “OR” took the logical sum of the geodesics passing at the two separated regions (e.g., hot spots or cold regions, defined below), each of which included a population of nodes, while “AND” took the logical product.

## Results

### Geodesics were dense nearby endpoint of fissures and sulci and sparse at cortical poles

To investigate the spatial distribution of the geodesics on the cortex (at 27 GW), we first observed the passage density of the geodesics at each position (See Materials and Methods). We found several regions exhibiting 100% higher density than the average over the cortical surface (Fig. 2A, red), while other regions exhibited 30% lower density than the average (Fig. 2A, blue). In the lateral view of the brain, high local density was observed in the perisylvian regions (red and yellow in Fig. 2A1), while low density was observed around the frontal, temporal, and occipital poles (blue in Fig. 2A1). Additionally, in the medial view, high local density emerged in the cingulate (red region around white-masked region in Fig. 2A2) and hippocampal gyri in the medial view (red in Fig. 2A2) and in the insula in the frontal view (red in Fig. 2A3). Hereafter, the regions with high and low local density are referred to as hot spots (HSs; Fig. 2A, red) and cold regions (CRs; Fig. 2A, blue), respectively. The HSs were surrounded by moderately high-density regions. In summary, the distribution of geodesics on the fetal cortex depended on the anatomical regions.

**Fig. 2:**
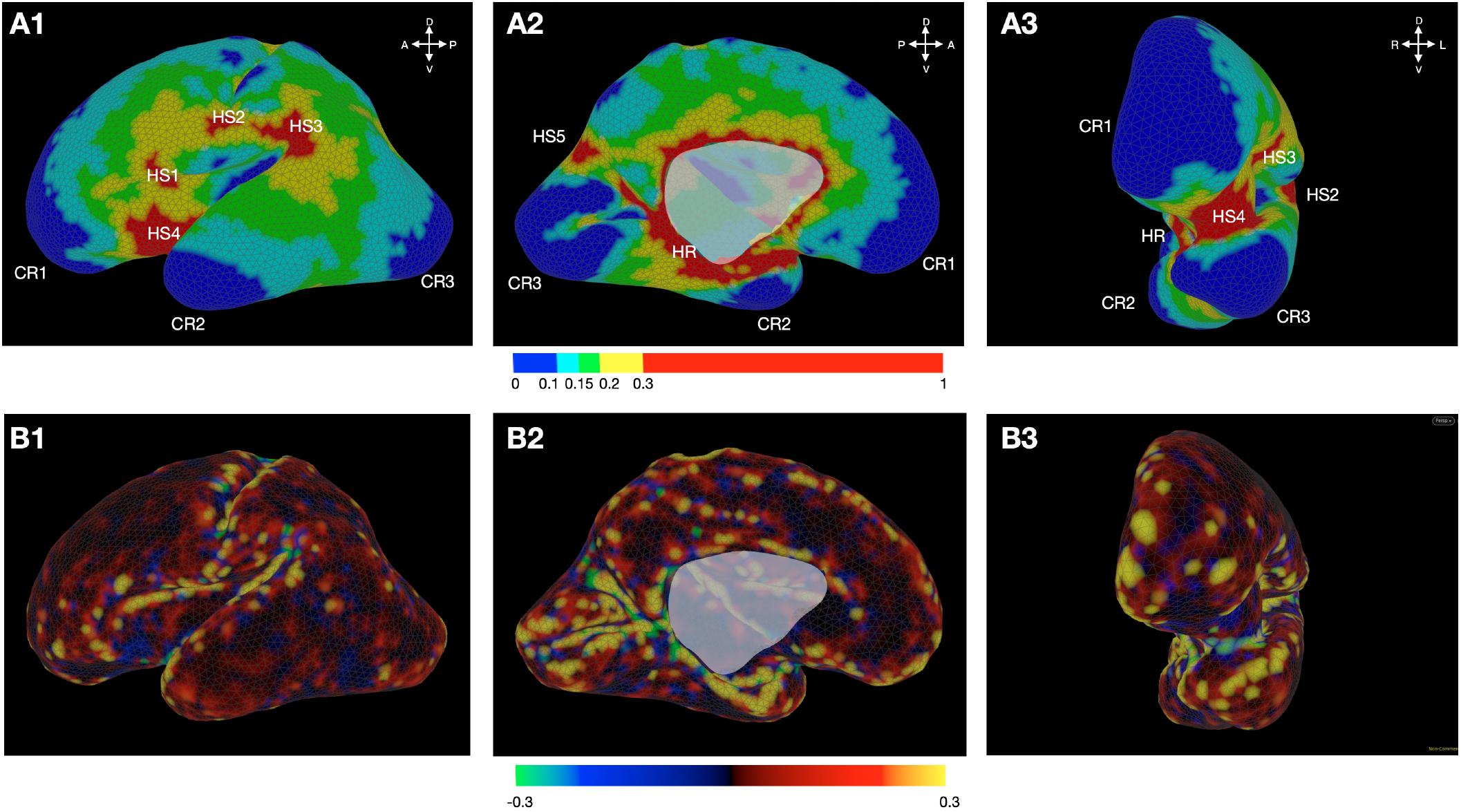
Association of geodesic distribution with cortical surface geometry. Heat map for normalized density of the geodesics (A1-3) and Gaussian curvature (B1-3) on the brain surface at 27 GW, where the inset of each panel denotes the respective value. Reference orientations: anterior (A), posterior (P), dorsal (D), ventral (V), left (L), and right (R) are placed in the right upper corner of each panel.

We found five HSs (hereafter, referred to as HS1, 2, 3, 4, and 5), each with a specific spatial distribution (Fig. 2A1 and 2, HS1-5). In the lateral view, HS1 was located around the inferior part of the frontal cortex (Fig. 2A1, HS1). HS2 was located around the anteroventral endpoint of the central sulcus and expanded posteriorly on the parietal cortex and anteriorly on the frontal cortex (Fig. 2A1, HS2).

HS3 was located around the endpoint of the Sylvian fissure (Fig. 2A1) and distributed anteriorly toward HS2 on the parietal cortex and ventrally on the temporal cortex (Fig. 2A1, HS3). In the lateral and frontal views, HS4 was located around the anterior part of the large intracortical insula (Fig. 2A1 and 3, HS4). In the medial view, HS5 was located around the parieto-occipital sulcus (Fig. 2A2, HS5). In addition, geodesics were densely distributed in a ring shape (hereafter referred to as Hot Ring, HR) in the cingulate and hippocampal gyri (Fig. 2A2, HR).

Moreover, we observed three CRs (hereafter referred to as CR1, 2, and 3) corresponding to the brain poles. CR1 was located at the frontal pole and expanded dorsally on the frontal cortex (Fig. 2A, CR1). CR2 was located at the temporal pole and expanded posteriorly on the temporal lobe (Fig. 2A, CR2). CR3 was located at the occipital pole and surrounded the calcarine sulcus in the medial view (Fig. 2A, CR3). To understand how the geodesic density relates to the local geometry, we measured the Gaussian curvature. The curvature on all five HSs and part of the HR tended to be negative, indicating a saddle shape at the endpoint of the gyri and sulci (Fig. 2B in blue or green). In contrast, all three CRs mostly had positive Gaussian curvature, exhibiting a dome-shape (Fig. 2B in red or yellow). These findings indicate that the typical geometrical landscape of the cortical surface biases the geodesic density.

### Dense geodesics corresponded to the position of two association fibers

The correspondence of the HSs and HR to the anatomical regions (Fig. 2A) prompted us to examine whether the geodesics further corresponded to major association tracts, such as the superior longitudinal fasciculus (slf)/arcuate fasciculus and cg, in reference to the MRI atlas of the white matter of the human brain (Oishi et al., 2011; Catani et al., 2012).

First, the geodesics passing through the HSs on the lateral surface were delineated to compare their distribution with the slf or arcuate fasciculus, a lateral tract composed of long and short fibers connecting the perisylvian cortex to the frontal, parietal, and temporal lobes. We separately selected the geodesics passing through both HS1 (right gray cubes, Fig. 3A) and HS2 (central gray cubes, Fig. 3A), as well as the geodesics passing through HS2 and HS3 (left gray cubes, Fig. 3A) using the “AND” operation. By further applying the “OR” operation to the geodesics passing through both HS1 and HS2 and those passing through both HS2 and HS3, we obtained a bundle of geodesics passing through HS1, 2, and 3 (See Materials and Methods). We found that this geodesic bundle traveled along the anterior-posterior direction over the moderately high-density region (Fig. 2A, yellow) of the frontal and parietal regions (white lines, Fig. 3A). Moreover, many geodesics passing through HS3 curved towards the temporal lobe, forming a large arc on the perisylvian cortex as shown in Fig. 3A. Thus, the shape and trajectory of the obtained bundle correspond to those of the slf or arcuate fasciculus.

**Fig. 3:**
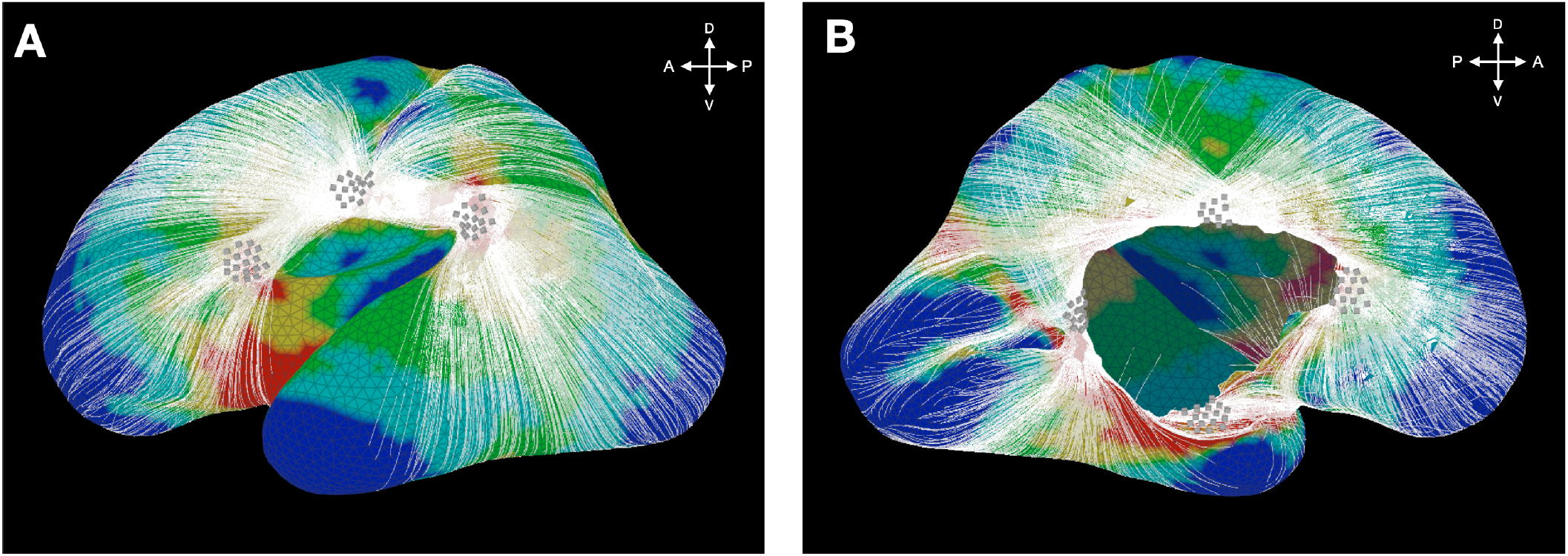
Dense geodesics. White line: geodesic. (A) The geodesic bundle obtained by applying the “OR” operation to two bundles selected by the “AND” operation passing through HS1 and HS2 and through HS2 and HS3. (B) The geodesic bundle obtained by applying the “OR” operation to three bundles selected by the “AND” operation passing through the anterior-dorsal part, dorsal-posterior part, and posterior-ventral part of the HR (ventral part: parahippocampal gyrus, dorsal part: isthmus cingulate gyrus, anterior part: anterior cingulate gyrus, posterior part: posterior cingulate gyrus). The colours of the heat map and reference orientations are the same as those in Fig. 2A.

Second, the geodesics passing through the HR on the medial surface were delineated to compare their distribution with the cg, a medial tract running immediately dorsal to the corpus callosum and along the ventral face of the hippocampus, forming a large, C-shaped trajectory (Oishi et al., 2011). We selected four parts of the HR: the anterior part on the anterior cingulate gyrus (right gray cubes, Fig. 3B), the dorsal part on the isthmus cingulate gyrus (upper gray cubes, Fig. 3B), the posterior part on the posterior cingulate gyrus (right gray cubes, Fig. 3B), and the ventral part on the parahippocampal gyrus (lower gray cubes, Fig. 3B). We applied the “OR” operation to three bundles of geodesics: those passing the anterior & dorsal parts, those passing the dorsal & posterior parts, and those passing the posterior & ventral parts, which were selected by the “AND” operation. As a result, a circular bundle of geodesics passing through the anterior cingulate, isthmus cingulate, posterior cingulate, and parahippocampal gyrus was obtained as shown in Fig. 3B. Moreover, a part of the bundle extended toward HS5 along the parieto-occipital fissure and a hot region along the calcarine fissure (Fig. 3B and Fig. 2A2). Thus, the shape of the obtained bundle of geodesics corresponded to that of the cg. In summary, dense geodesics passing through HSs and HR correspond to the position of slf and cg association fibers (Table. 1).

**Table 1.** Geodesic bundles passing through HSs and CRs corresponded to major association fibers

| Geodesic bundles | Geodesic characteristics | Association fiber |
| --- | --- | --- |
| HS1 - HS2 - HS3 | High density | Superior longitudinal fasciculus (slf) / Arcuate fasciculus |
| HR - HS5 | High density | Cingulum (cg) |
| CR1 - HS4 - CR2 | Inter-poles | Uncinate fasciculus (unc) |
| CR1 - HS4 - CR3 | Inter-poles | Inferior fronto-occipital fasciculus (ifo) |
| CR2 - HR - CR3 | Inter-poles | Inferior longitudinal fasciculus (ilf) |

### Interconnected geodesics between brain poles corresponded to the other association fibers

We also observed geodesic bundles interconnected between brain poles (hereafter referred to as inter-pole geodesics), despite the CRs (Fig. 4, CR1, CR2, and CR3) at the cortical poles. We examined whether the inter-pole geodesics corresponded to the fronto-temporal (unc), fronto-occipital (ifo), and temporo-occipital (inferior longitudinal fasciculi (ilf)) inter-pole association fibers, according to the MRI atlas (Oishi et al., 2011; Catani et al., 2012).

**Fig. 4:**
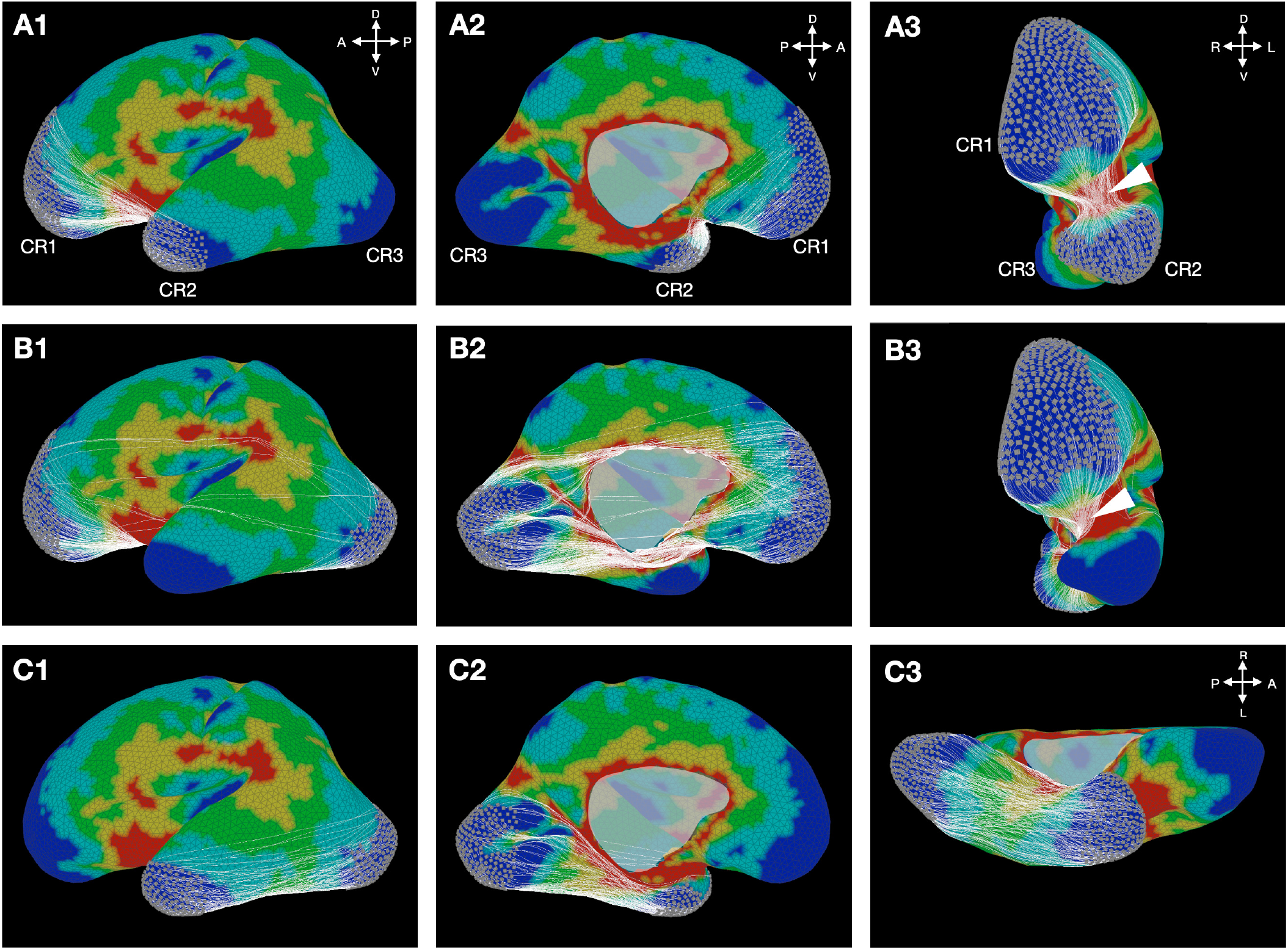
Inter-pole geodesics connecting brain poles. The bundle of geodesics with start and end positions located within CR1 and CR2 (A); CR1 and CR3 (B); and CR2 and CR3 (C). The white arrow points to the position where the bundle passes through HS4. The colours of the heat map and reference orientations are the same as those in Fig. 2A.

The inter-pole geodesics connecting the inferior part of the frontal cortex (CR1) and the temporal pole (CR2) exhibited a converged passage through a narrow part of HS4 (Fig.4A3). This bundle of geodesics corresponded to the unc, a tract that connects the medial and lateral orbitofrontal cortex with the anterior temporal lobe and passes through the narrow region of the external capsule (white arrow, Fig. 4A3).

The inter-pole geodesics connecting CR1 and the inferior and medial occipital cortex (CR3) exhibited a converged passage through a narrow part of HS4 and the hot regions on the medial occipito-temporal cortex (Fig. 4B). This bundle corresponded to the inferior fronto-occipital fasciculus, a tract that connects the orbitofrontal cortex and the inferior and medial occipital lobes passing through the narrow region of the external capsule (Fig. 4B2 and white arrow, 4B3). Furthermore, the tangential relative positions of the geodesic bundles (compare white arrow between Fig. 4B3 and 4A3) were consistent with the anatomical property in which the fibers of the ifo enter the external capsule dorsally to the fibers of the unc.

The inter-pole geodesics between CR2 and CR3 passed through the ventro-medial HR on the occipito-temporal cortex, as shown in Fig. 4C in yellow. This bundle corresponded to the ilf, a tract connecting the occipital and temporal lobes. Moreover, the tangential relative position of this bundle with respect to the bundle between CR1 and CR3 (compare Fig. 4C1 with Fig. 4B1) was consistent with the anatomical property in which the ilf runs parallel and lateral to the ifo. In summary, the inter-pole geodesics for fronto-temporal, fronto-occipital, and temporo-occipital connectivity corresponded to the position of inter-pole association fibers (Table.1).

## Discussion

### Summary of results

As a hypothetical landmark for tangential long-distance fiber guidance, we calculated the spatial distribution of the geodesics on the fetal cortical surface, where association fiber prototypes have already emerged (Horgos et al., 2020; Takahashi et al., 2012; Huang et al., 2009). We found that geodesics were dense at the saddle-shaped anatomical regions (HSs and HR in Fig. 2A) and sparse at the dome-shaped cortical poles (CRs in Fig. 2B). The positions of the multiple geodesics passing through the HSs and HR corresponded to those of the two major association fibers (Fig. 3 and Table.1), while those connecting the CRs at the cortical poles via the HSs or HR corresponded to the three other major association fibers (Fig.3, 4, and Table.1). These correspondent spatial distributions between association fibers and geodesics support that geodesics guide the tangential position of the long-distance association fibers according to the curved cortical surface landscape. This geodesic theory expands upon the subplate nexus hypothesis to provide a better understanding of the directional guidance of long-distance association fibers.

### A hypothetical mechanism for the development of association fibers

Neuronal activity serves as a potential physiological mechanism for the geodesic-mediated directional guidance. Subplate neurons exhibit spontaneous activity forming neural circuits via gap junctions and chemical synapses (Moore et al., 2011, 2014). These neuronal circuits have been shown to play an important role in tangential neuronal fibers (Kostović, 2020), as well as in radial fibers (Kanold and Luhmann, 2010; Kanold, 2019; Molnár et al., 2020, 2019; Luhmann et al., 2022; Bystron et al., 2008; Hadders-Algra, 2018; Ohtaka-Maruyama, 2020). In the tangential direction, neuronal activity can be propagated to the subplate neural circuits throughout the entire brain (Thomason et al., 2017; Keunen et al., 2017; Kilb et al., 2011; Garaschuk et al., 2000; Sun and Luhmann, 2007). A theoretical study has shown that such propagation of neuronal excitatory activity follows a geodesics on the curved surface of propagated media (e.g., brain surface) (Horibe et al., 2019; Davydov et al., 2000; Brazhnik et al., 1988). This suggests that neuronal activity can be densely propagated at the HSs (Fig.1 and 4), thereby guiding neuronal fibers in a similar way to Hebbian learning (Hebb, 1949). The network of subplate neurons is connected to the cortical neurons, which migrate in the radial direction towards the surface (Ohtaka-Maruyama et al., 2018). Cortical neurons grow their tangential network, using the network of subplate neurons as the prototype. When the subplate disappears via apoptosis during the fetal period (Kostovic and Rakic, 1990), the network of subplate neurons is replaced by corticocortical networks (i.e., association fibers networks) (Kostović, 2020). Together, the previous and present studies suggest that the association fibers network is formed by tangential propagation of neuronal activities following geodesics on the smooth surface of the fetal brain.

### The geodesic theory is applicable to other developmental stages and species

The geodesic theory can be extended to other development stages. Smoothly curved landscapes, such as the saddle shape around the Sylvian fissure (Fig. 2A3, around HS4) and the dome shape at the brain poles (Fig. 2A1, around CR1-3) are maintained between 9 and 27 GW, despite the volumetric growth of the cortex (Horgos et al., 2020; Huang et al., 2009). For example, at 9 GW, primary folding has already begun to develop, resulting in a saddle-shaped Sylvian fissure, while frontal and temporal poles exhibit a dome shape (Horgos et al., 2020). This consistency in the surface landscape suggests that the HSs of geodesics on the saddles (Fig.2A, HS4) and the CRs on the domes interconnected by the geodesics (Fig.4A) exist even at 9 GW. Importantly, the unc appears to connect the brain poles around 9 GW, which is the earliest connection among the association fibers (Horgos et al., 2020; Ouyang et al., 2019). Thus, the position of association fiber formation could correspond to the geodesics earlier than 27 GW. At later stages, short association fibers (e.g., U-shaped fiber) emerge along with a shorter-distance landscape of the secondary sulci than that of primary sulci (Takahashi et al., 2012; Das and Takahashi, 2018). Thus, whether the position of short-distance fibers corresponds to the geodesics of the shorter-distance landscape should also be evaluated. In summary, examining the geodesic theory in early and late fetal developmental stages would provide directional guidance of long and short association fibers, respectively.

The geodesic theory is further applicable to other species. Primates share the primary sulci; however, species evolution modifies their brain surface landscape such as the depth and length of the sulci (Friedrich et al., 2021; Miller et al., 2020; Amiez et al., 2019). This morphological evolution of shared anatomical characteristics enables the modification of the respective distribution of geodesics, potentially affecting that of the association fibers. For example, the arcuate fasciculus is well-developed (strongly connected) in humans, playing an important role in language function (Dick and Tremblay, 2012); however, it is immaturely and weakly connected in other primates (Rilling et al., 2008). In the human fetal cortex, the spatial distribution of geodesics corresponded to those of the arcuate fasciculus (HS1, HS2, HS3; Fig. 2A and Fig. 3A). Future studies should examine whether and how the density and direction of geodesics correspond to those of the arcuate fasciculus and other association fibers in other primate cortices, which would provide further understanding of the evolution of association fibers constrained by the cortical landscape.

### Future implications

The present theory does not incorporate the radial thickness of the subplate depending on the growth stages and cortical regions (Terashima et al., 2021; Huang et al., 2009; Garcia et al., 2018a). Nevertheless, the geodesic bundles corresponded to the five major association fibers. The effect of the thickness on the geodesic distribution can be evaluated if the sampling probability of the start and end positions of the geodesics is set to depend on the respective thickness; however, in this study, this probability was set to be homogeneous. Additionally, the geodesic theory includes the cortical surface geometry but not the radial direction. Indeed, the ifo connecting fronto-occipital poles (Fig.4B and 4C) and the ilf connecting temporo-occipital poles (Fig.4B and 4C) are interiorly positioned from the cortical surface. Despite the absence of depth information, the geodesics reproduced the relative tangential position of the fibers. For example, the inter-pole geodesics between fronto-occipital poles corresponding to the ifo were located more dorsally than those between fronto-temporal poles corresponding to the unc (Fig.4A3 and 4B3 white arrows), consistent with the fact that the ifo passes through a more dorsal area than the unc around the external capsule (Catani et al., 2012). Furthermore, the geodesics corresponding to the ilf and ifo reproduce this tangential relative position, in which the ilf runs parallel to the outer part of the ifo (Catani et al., 2012) (Fig.4B2 and 4C2). The reliability of the geodesic theory on the tangential fiber distribution should be further evaluated by integrating diffusion tensor imaging (DTI) tract data with the geodesic position data.

### Conclusion

Our geometrical analysis revealed that the distribution of geodesics on the cortical surface of human fetal brains follows that of the five major association fibers. Therefore, we proposed and confirmed the geodesic theory, in which the tangential direction of the association fibers follows geodesics, suggesting a unified understanding on the development and evolution of association fibers constrained by the landscape of the cortical surface.

## Notes

We thank K. Matsushita for discussion and J. Horikawa for technical support in Houdini.

## Author contributions statement

KH, GT, and KF conceived the experiments, KH conducted the experiments, KH, GT and KF analysed the results. KH, GT and KF wrote and reviewed the manuscript.

## References

Amiez, C., Sallet, J., Hopkins, W. D., Meguerditchian, A., Hadj-Bouziane, F., Ben Hamed, S., Wilson, C. R. E., Procyk, E., and Petrides, M. (2019). Sulcal organization in the medial frontal cortex provides insights into primate brain evolution. Nature Communications, 10(1):3437.

Brazhnik, P. K., Davydov, V. A., and Mikhailov, A. S. (1988). Kinematic approach to the description of autowave processes in active media. Theor. Mat. Fiz., 74:440–447.

Bystron, I., Blakemore, C., and Rakic, P. (2008). Development of the human cerebral cortex: Boulder Committee revisited. Nature Reviews Neuroscience, 9(2):110–122.

Catani, M., Schotten, D., and Thiebaut, M. (2012). Atlas of human brain connections. Oxford University Press.

Comer, J. D., Alvarez, S., Butler, S. J., and Kaltschmidt, J. A. (2019). Commissural axon guidance in the developing spinal cord: from Cajal to the present day. Neural Development, 14(1):9.

Das, A. and Takahashi, E. (2018). Neuronal migration and axonal pathways linked to human fetal insular development revealed by diffusion MR tractography. Cerebral Cortex, 28(10):3555–3563.

Davydov, V., Morozov, V., and Davydov, N. (2000). Ring-shaped autowaves on curved surfaces. Physics Letters A, 267(5-6):326–330.

Dick, A. S. and Tremblay, P. (2012). Beyond the arcuate fasciculus: Consensus and controversy in the connectional anatomy of language. Brain, 135(12):3529–3550.

Duque, A., Krsnik, Z., Kostović, I., and Rakic, P. (2016). Secondary expansion of the transient subplate zone in the developing cerebrum of human and nonhuman primates. Proceedings of the National Academy of Sciences of the United States of America, 113(35):9892–9897.

Egger, K., Yang, S., Reisert, M., Kaller, C., Mader, I., Beume, L., Weiller, C., and Urbach, H. (2015). Tractography of Association Fibers Associated with Language Processing. Clinical Neuroradiology, 25:231–236.

Friedrich, P., Forkel, S. J., Amiez, C., Balsters, J. H., Coulon, O., Fan, L., Goulas, A., Hadj-Bouziane, F., Hecht, E. E., Heuer, K., et al. (2021). Imaging evolution of the primate brain: the next frontier? NeuroImage, 228:117685.

Garaschuk, O., Linn, J., Eilers, J., and Konnerth, A. (2000). Large-scale oscillatory calcium waves in the immature cortex. Nature Neuroscience, 3(5):452–459.

Garcia, K. E., Kroenke, C. D., and Bayly, P. V. (2018a). Mechanics of cortical folding: stress, growth and stability. Philosophical Transactions of the Royal Society B: Biological Sciences, 373(1759):20170321.

Garcia, K. E., Robinson, E. C., Alexopoulos, D., Dierker, D. L., Glasser, M. F., Coalson, T. S., Ortinau, C. M., Rueckert, D., Taber, L. A., Van Essen, D. C., Rogers, C. E., Smyser, C. D., and Bayly, P. V. (2018b). Dynamic patterns of cortical expansion during folding of the preterm human brain. Proceedings of the National Academy of Sciences, 115(12):3156–3161.

Hadders-Algra, M. (2018). Early human brain development: Starring the subplate. Neuroscience & Biobehavioral Reviews, 92:276–290.

Hebb, D. (1949). The Organization of Behavior. New York: Wiley & Sons.

Horgos, B., Mecea, M., Boer, A., Szabo, B., Buruiana, A., Stamatian, F., Mihu, C.-M., Florian, I. Ş., Susman, S., and Pascalau, R. (2020). White matter dissection of the fetal brain. Frontiers in Neuroanatomy, 14:584266.

Horibe, K., Hironaka, K.-i., Matsushita, K., and Fujimoto, K. (2019). Curved surface geometry-induced topological change of an excitable planar wavefront. Chaos: An Interdisciplinary Journal of Nonlinear Science, 29(9):093120.

Huang, H., Xue, R., Zhang, J., Ren, T., Richards, L. J., Yarowsky, P., Miller, M. I., and Mori, S. (2009). Anatomical characterization of human fetal brain development with diffusion tensor magnetic resonance imaging. Journal of Neuroscience, 29(13):4263–4273.

Kanold, P. O. (2019). The first cortical circuits: Subplate neurons lead the way and shape cortical organization. Neuroforum, 25(1):15–24.

Kanold, P. O. and Luhmann, H. J. (2010). The Subplate and Early Cortical Circuits. Annual Review of Neuroscience, 33(1):23–48.

Keunen, K., Counsell, S. J., and Benders, M. J. (2017). The emergence of functional architecture during early brain development. NeuroImage, 160:2–14.

Kilb, W., Kirischuk, S., and Luhmann, H. J. (2011). Electrical activity patterns and the functional maturation of the neocortex. European Journal of Neuroscience, 34(10):1677–1686.

Kimmel, R. and Sethian, J. A. (1998). Computing geodesic paths on manifolds. Proceedings of the National Academy of Sciences of the United States of America, 95(15):8431–8435.

Kobayashi, Shoshichi and Nomizu, K. (1963). Foundations of differential geometry. New York, London.

Kostović, I. (2020). The enigmatic fetal subplate compartment forms an early tangential cortical nexus and provides the framework for construction of cortical connectivity. Progress in Neurobiology, 194:101883.

Kostović, I., Išasegi, I. Ž., and Krsnik, Ž. (2019). Sublaminar organization of the human subplate: developmental changes in the distribution of neurons, glia, growing axons and extracellular matrix. Journal of anatomy, 235(3):481–506.

Kostovic, I. and Rakic, P. (1990). Developmental history of the transient subplate zone in the visual and somatosensory cortex of the macaque monkey and human brain. Journal of Comparative Neurology, 297(3):441–470.

Kostović, I., Sedmak, G., and Judaš, M. (2019). Neural histology and neurogenesis of the human fetal and infant brain. Neuroimage, 188:743–773.

Kostović, I., Seress, L., Mrzljak, L., and Judaš, M. (1989). Early onset of synapse formation in the human hippocampus: A correlation with Nissl-Golgi architectonics in 15-and 16.5-week-old fetuses. Neuroscience, 30(1):105–116.

Kristt, D. A. and Molliver, M. E. (1976). Synapses in newborn rat cerebral cortex: a quantitative ultrastructural study. Brain Research, 108(1):180–186.

Luhmann, H. J., Kanold, P. O., Molnár, Z., and Vanhatalo, S. (2022). Early brain activity: translations between bedside and laboratory. Progress in neurobiology, page 102268.

Luhmann, H. J. and Khazipov, R. (2018). Neuronal activity patterns in the developing barrel cortex. Neuroscience, 368:256–267.

Miller, J. A., Voorhies, W. I., Li, X., Raghuram, I., Palomero-Gallagher, N., Zilles, K., Sherwood, C. C., Hopkins, W. D., and Weiner, K. S. (2020). Sulcal morphology of ventral temporal cortex is shared between humans and other hominoids. Scientific Reports, 10(1):17132.

Molnár, Z., Clowry, G. J., Šestan, N., Alzu’bi, A., Bakken, T., Hevner, R. F., Hüppi, P. S., Kostović, I., Rakic, P., Anton, E., et al. (2019). New insights into the development of the human cerebral cortex. Journal of anatomy, 235(3):432–451.

Molnár, Z., Luhmann, H. J., and Kanold, P. O. (2020). Transient cortical circuits match spontaneous and sensory-driven activity during development. Science, 370(6514):eabb2153.

Moore, A. R., Zhou, W.-L., Jakovcevski, I., Zecevic, N., and Antic, S. D. (2011). Spontaneous Electrical Activity in the Human Fetal Cortex In Vitro. Journal of Neuroscience, 31(7):2391–2398.

Moore, A. R., Zhou, W. L., Sirois, C. L., Belinsky, G. S., Zecevic, N., and Antic, S. D. (2014). Connexin hemichannels contribute to spontaneous electrical activity in the human fetal cortex. Proceedings of the National Academy of Sciences of the United States of America, 111(37):E3919–E3928.

Mori, S., Wakana, S., Van Zijl, P. C., and Nagae-Poetscher, L. (2005). MRI Atlas of Human White Matter. Elsevier.

Ohtaka-Maruyama, C. (2020). Subplate neurons as an organizer of mammalian neocortical development. Frontiers in Neuroanatomy, 14:8.

Ohtaka-Maruyama, C., Okamoto, M., Endo, K., Oshima, M., Kaneko, N., Yura, K., Okado, H., Miyata, T., and Maeda, N. (2018). Synaptic transmission from subplate neurons controls radial migration of neocortical neurons. Science, 360(6386):313–317.

Oishi, K., Mori, S., Donohue, P. K., Ernst, T., Anderson, L., Buchthal, S., Faria, A., Jiang, H., Li, X., Miller, M. I., van Zijl, P. C., and Chang, L. (2011). Multi-contrast human neonatal brain atlas: Application to normal neonate development analysis. NeuroImage, 56(1):8–20.

Ouyang, M., Dubois, J., Yu, Q., Mukherjee, P., and Huang, H. (2019). Delineation of early brain development from fetuses to infants with diffusion mri and beyond. Neuroimage, 185:836–850.

Philippi, C. L., Mehta, S., Grabowski, T., Adolphs, R., and Rudrauf, D. (2009). Damage to association fiber tracts impairs recognition of the facial expression of emotion. Journal of Neuroscience, 29(48):15089–15099.

Rilling, J. K., Glasser, M. F., Preuss, T. M., Ma, X., Zhao, T., Hu, X., and Behrens, T. E. J. (2008). The evolution of the arcuate fasciculus revealed with comparative DTI. Nature Neuroscience, 11(4):426–428.

Sethian, J. A. (1996). A fast marching level set method for monotonically advancing fronts. Proceedings of the National Academy of Sciences of the United States of America, 93(4):1591–1595.

Stoeckli, E. T. (2018). Understanding axon guidance: are we nearly there yet? Development, 145(10):dev151415.

Sun, J. J. and Luhmann, H. J. (2007). Spatio-temporal dynamics of oscillatory network activity in the neonatal mouse cerebral cortex. European Journal of Neuroscience, 26(7):1995–2004.

Takahashi, E., Folkerth, R. D., Galaburda, A. M., and Grant, P. E. (2012). Emerging cerebral connectivity in the human fetal Brain: An MR tractography study. Cerebral Cortex, 22(2):455–464.

Terashima, M., Ishikawa, A., Mánner, J., Yamada, S., and Takakuwa, T. (2021). Early development of the cortical layers in the human brain. Journal of Anatomy, 239(5):1039–1049.

Thomason, M. E., Scheinost, D., Manning, J. H., Grove, L. E., Hect, J., Marshall, N., Hernandez-Andrade, E., Berman, S., Pappas, A., Yeo, L., Hassan, S. S., Constable, R. T., Ment, L. R., and Romero, R. (2017). Weak functional connectivity in the human fetal brain prior to preterm birth. Scientific Reports, 7:1–10.

Van Essen, D. C. (2020). A 2020 view of tension-based cortical morphogenesis. Proceedings of the National Academy of Sciences, 117(52):32868–32879.

Vasung, L., Lepage, C., Radoš, M., Pletikos, M., Goldman, J. S., Richiardi, J., Raguž, M., Fischi-Gómez, E., Karama, S., Huppi, P. S., et al. (2016). Quantitative and qualitative analysis of transient fetal compartments during prenatal human brain development. Frontiers in neuroanatomy, 10:11.

Vasung, L., Turk, E. A., Ferradal, S. L., Sutin, J., Stout, J. N., Ahtam, B., Lin, P.-Y., and Grant, P. E. (2019). Exploring early human brain development with structural and physiological neuroimaging. Neuroimage, 187:226–254.

